# The CUL4-DDB1-DCAF1 E3 ubiquitin ligase complex regulates PLK4 protein levels to prevent premature centriole duplication

**DOI:** 10.1101/2023.09.13.555514

**Authors:** Josina Grossmann, Anne-Sophie Kratz, Alina Kordonsky, Gali Prag, Ingrid Hoffmann

## Abstract

Centrioles play important roles in the assembly of centrosomes and cilia. Centriole duplication occurs once per cell cycle and is dependent on polo-like kinase 4 (PLK4). To prevent centriole amplification, which is a hallmark of cancer, PLK4 protein levels need to be tightly regulated. Here, we show that the Cullin(CUL)4A/B-DDB1-DCAF1, CRL4^DCAF1^, E3 ligase targets PLK4 for degradation in human cells. DCAF1 binds and ubiquitylates PLK4 in G2 phase to prevent premature centriole duplication in mitosis. In contrast to the regulation of PLK4 by SCF^β-TrCP^, the interaction between PLK4 and DCAF1 is independent of PLK4 kinase activity and mediated by polo-boxes 1 and 2 of PLK4, suggesting that DCAF1 promotes PLK4 ubiquitylation independently of β-TrCP. Thus, the SCF^Slimb/β-TrCP^ pathway, targeting PLK4 for ubiquitylation based on its phosphorylation state and CRL4^DCAF1^, which ubiquitylates PLK4 by binding to the conserved PB1-PB2 domain, appear to be complementary ways to control PLK4 abundance to prevent centriole overduplication.

## Introduction

Centriole biogenesis has to be tightly controlled to prevent aberrant centrosome number, which can lead to chromosome missegregation and aneuploidy and has been associated with cancer (Nigg and Holland, 2018). Centriole duplication is triggered by and dependent on polo-like kinase 4 (PLK4), a divergent member of the polo-like kinase family (Bettencourt-Dias et al., 2005; Habedanck et al., 2005). Binding of PLK4 to its centriole substrate STIL promotes activation of the kinase (Ohta et al., 2014; Moyer et al., 2015; Lopes et al., 2015). PLK4 phosphorylates STIL in a conserved STAN motif which leads to binding and recruitment of SAS6 (Ohta et al., 2014; Dzhindzhev et al., 2014; Moyer et al., 2015; Kratz et al., 2015), which is necessary for cartwheel assembly (Nakazawa et al., 2007).

PLK4 protein levels are regulated by ubiquitylation and proteasomal degradation. Previous work has revealed that this is in part mediated by the SCF (Skp1/Cullin/F-box) -β-TrCP/Slimb E3 ubiquitin ligase (Rogers et al., 2009; Cunha-Ferreira et al., 2009; Guderian et al., 2010). The β−propeller of the F-box protein β-TrCP recognizes a conserved phosphodegron in the N-terminal PEST motif of PLK4, which is generated by homodimer-dependent trans-autophosphorylation of human PLK4 (Guderian et al., 2010; Holland et al., 2010; Cunha-Ferreira et al., 2013; Klebba et al., 2013).

CUL4-RING ligases contain the scaffold proteins CUL4A or CUL4B, which are conserved from yeast to humans. They bind to a substrate-targeting unit, which is composed of the adaptor DDB1 and a member of the DDB1- and CUL4-associated factors (DCAF), a family of WD40 repeat proteins that confer substrate specificity (Jackson and Xiong, 2009). Among the DCAFs, DCAF1 is a critical substrate receptor in the DCAF1-DDB1-CUL4 complex (Han et al., 2020). DCAF1 is also known as Vpr binding protein (VprBP), as it was initially discovered as a target protein hijacked by the viral protein Vpr of human immunodeficiency virus-1 (HIV-1) (Tan et al., 2007). DCAF1 is involved in a number of fundamental cellular processes including DNA replication (McCall et al., 2008) and cell cycle regulation (Guo et al., 2016). A number of substrates of the CUL4-DDB1-DCAF1 complex CRL4^DCAF1^ have been described. Among those are p53 (Hrecka et al., 2007; Guo et al., 2016), the replication factor MCM10 (Kaur et al., 2012) and protein phosphatase 2A (Yu et al., 2015). The activity of the CUL4-DDB1-DCAF1 complex itself is regulated by its oligomerization state (Mohamed et al., 2021).

Recent data showed that a β-TrCP binding mutant of PLK4 was still ubiquitylated and only modestly stabilized in human cells, suggesting that additional ubiquitin ligases might regulate PLK4 protein levels in canonical centriole duplication (Rogers et al., 2009; Holland et al., 2010; Klebba et al., 2013). Here, we report that the CRL4^DCAF1^ E3 ubiquitin ligase complex contributes to the regulation of PLK4 abundance. DCAF1 binds to and promotes ubiquitylation of PLK4. Alphafold2.0 modelling corroborated by *in vivo* analysis demonstrates a novel binding interface in which the unstructured DCAF1 acidic tail binds to the groove of the PLK4 polo-boxes 1 and 2. Furthermore, we find that the CRL4^DCAF1^ complex controls PLK4 levels in G2 phase, when β-TrCP activity is low (Paul et al., 2022), thus preventing premature centriole disengagement and centriole duplication. The interaction between DCAF1 and PLK4 and the ubiquitylation of PLK4 occur in a PLK4 kinase activity independent and phosphorylation independent manner, suggesting that the SCF^β-TrCP^ and the CRL4^DCAF1^ E3 ubiquitin ligases independently control PLK4 protein abundance and thereby centriole duplication.

## Results

### 1. CUL4-DDB1-DCAF1 E3 ubiquitin ligase, CRL4^DCAF1^, regulates PLK4 protein levels

To identify ubiquitin ligases that regulate PLK4 protein levels, we performed co-immunoprecipitation experiments with PLK4 as bait followed by subsequent mass spectrometry analysis (Fig. S1A). Interestingly, amongst other known PLK4-interacting proteins such as STIL, CEP152 and β-TrCP, we identified the substrate recognition component DCAF1 (VprBP), along with DNA-damage binding protein 1 (DDB1), a core component of CUL4A and CUL4B-based E3 ubiquitin ligases (Jackson and Xiong, 2009). We confirmed the interaction between overexpressed PLK4 and endogenous DCAF1 (Fig. 1A). In addition, an interaction between endogenous PLK4 and DCAF1 was observed by using specific antibodies (Fig. 1B). Apart from its interaction with DCAF1, we found that PLK4 also interacted with DDB1, CUL4A and CUL4B after co-expression (Fig. S1B). Sequential co-immunoprecipitation experiments revealed that PLK4 exists in a complex with DCAF1, DDB1 and CUL4 but not with EDD, a HECT-type E3 ubiquitin ligase which also contains DCAF1 as substrate-binding domain (Fig. S1C) (Maddika and Chen, 2009). DCAF1 is a centrosome protein (Hossain et al., 2017) and DDB1 was previously identified in a proteomic approach to define the constituents of human centrosomes (Andersen et al., 2003). Autophosphorylation of two amino acid residues (S285/T289 in human PLK4) within the PLK4 β-TrCP-binding motif promotes the binding of β-TrCP and subsequent ubiquitylation and destruction of PLK4 (Guderian et al., 2010; Holland et al., 2010; Cunha-Ferreira et al., 2013; Klebba et al., 2013). We first aimed at determining whether DCAF1 would bind to the same or a different site within PLK4 than β-TrCP and whether this binding is dependent on the autophosphorylation of PLK4. To address this question, we used several mutants of PLK4. First, we generated a kinase-dead version of PLK4 by a mutation within the kinase domain (K41R) (Bahtz et al., 2012), second, we mutated either the β-TrCP recognition motif in PLK4 to AA (S285A+T289A) or third, we deleted the PEST destruction motif (Fig. 1C). Whereas binding between PLK4 and β-TrCP is lost when PLK4 is kinase-dead or mutated in the recognition motif, DCAF1 interacted with the kinase-dead PLK4 mutant, the β-TrCP-binding mutant PLK4-AA and with the PLK4-ΔPEST mutant to a similar level as with wild-type PLK4 (Fig. 1D). Consistent with this, by using the indicated PLK4 fragments, we could clearly map the binding site of DCAF1 to a C-terminal fragment of PLK4 containing the tandem polo-boxes (PB1-PB2) (Fig. S2). To further prove that binding between PLK4 and DCAF1 is independent of PLK4 autophosphorylation or phosphorylation by any other unknown kinase, we treated immunoprecipitated PLK4 with λ-phosphatase and find that while binding to β-TrCP was reduced, no reduction of DCAF1 binding to PLK4 could be observed (Fig. 1E). Together, these data suggest that autophosphorylation of PLK4 or phosphorylation of PLK4 in general is not required for the interaction between DCAF1 and PLK4.

**Figure 1.**
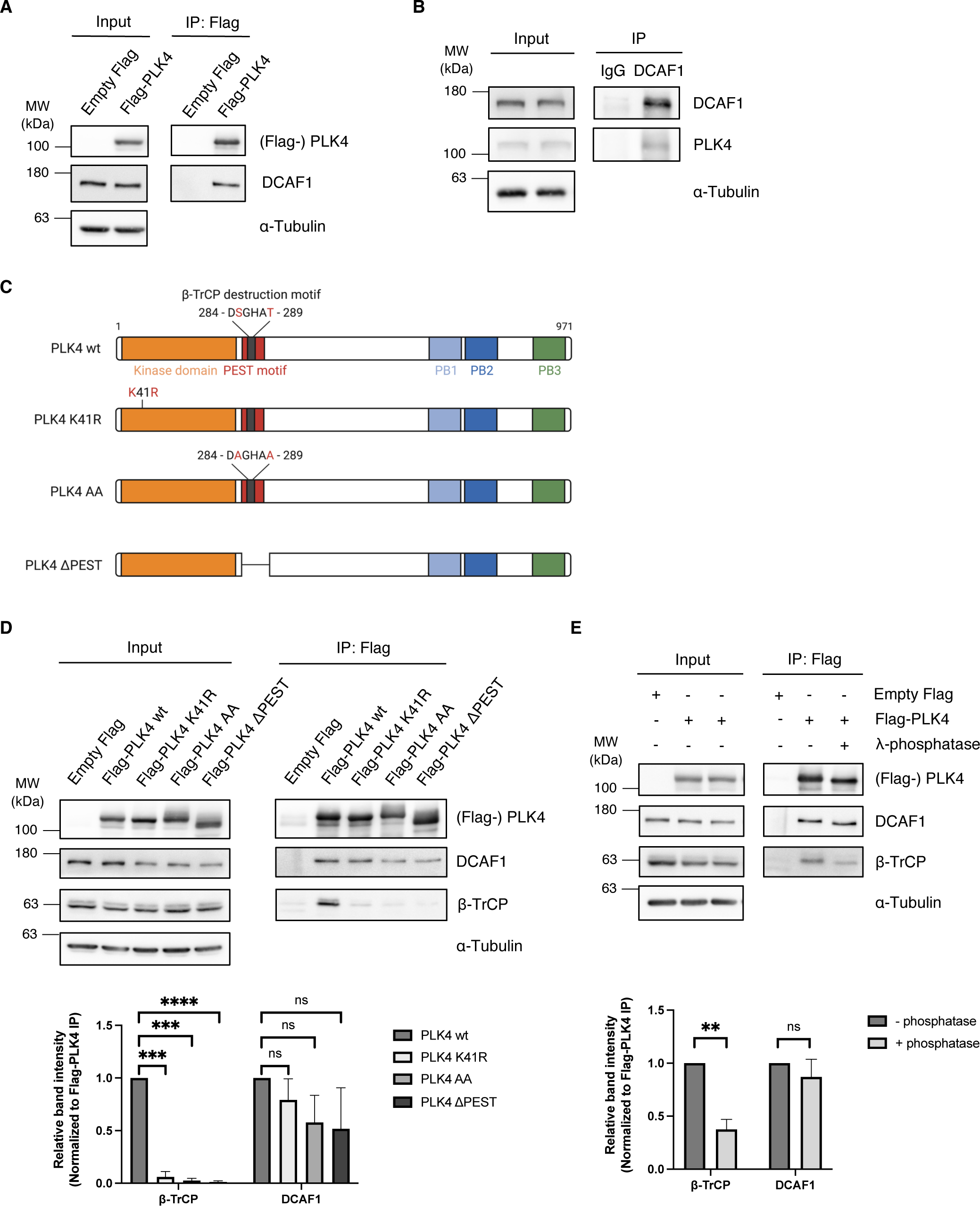
PLK4 interacts with DCAF1 independently of PLK4 kinase activity and phosphorylation. **(A)** Flag-PLK4 was overexpressed in HEK293T cells for 48 h. Co-precipitated DCAF1 was detected by IP against the Flag tag and subsequent Western blot analysis. **(B)** Endogenous DCAF1 was immunoprecipitated from HEK293T cell lysates using unspecific IgG control or specific DCAF1 antibodies and protein G sepharose. **(C)** Overview of PLK4 mutants used in (D). **(D)** Flag-PLK4 wt or Flag-PLK4 mutants (K41R, AA, ΔPEST) were overexpressed in HEK293T cells for 48 h. Co-precipitated DCAF1 and β-TrCP were detected by IP against the Flag tag and subsequent Western blot analysis. Quantification of relative β-TrCP/ Flag-PLK4 or DCAF1/ Flag-PLK4 signal normalized to Flag-PLK4 wt. **** p < 0.0001, *** p < 0.001, ns p > 0.05, unpaired Student’s t test with Welch’s correction, N = 3. **(E)** Flag-PLK4 was overexpressed in HEK293T cells for 48 h. Co-precipitated DCAF1 and β-TrCP were detected by IP against the Flag tag with or without dephosphorylation of protein samples using λ-phosphatase and subsequent Western blot analysis. Quantification of relative β-TrCP/ Flag-PLK4 or DCAF1/ Flag-PLK4 signal. ** p < 0.01, ns p > 0.05, unpaired Student’s t test with Welch’s correction, N = 3.

We further investigated whether DCAF1 depletion would affect PLK4 protein turnover. Treatment of cells with DCAF1 specific siRNAs led to an increase in PLK4 protein levels (Fig. 2A). The increase in PLK4 protein levels was not detectable upon depletion of another DCAF family member, DCAF5 (Zhang et al., 2019), which was also detected in our screen (Fig. 2B and Fig. S1A). Treatment of DCAF1 depleted cells with cycloheximide (CHX) to block protein translation led to stabilization of PLK4 and a slight increase in protein half-life (Fig. 2C). The effect on PLK4 protein levels was also clearly visible when we depleted DCAF1 in a doxycycline-inducible HeLa cell line. The time-dependent decrease in DCAF1 protein levels resulted in an increase in PLK4 protein levels (Fig. 2D). Depletion of DCAF1 also led to an increase in PLK4 protein levels at the centrosome (Fig. 2E). We anticipate that increased PLK4 protein levels upon depletion of DCAF1 should trigger centriole overduplication (Bettencourt-Dias et al., 2005; Habedanck et al., 2005). To assess this hypothesis, we depleted DCAF1 by addition of doxycycline and observed supernumerary centrioles leading to the formation of multipolar spindles in mitosis (Fig. 2F). Our data suggest that CRL4^DCAF1^ might function to keep PLK4 protein levels low, thus preventing centriole overduplication.

**Figure 2.**
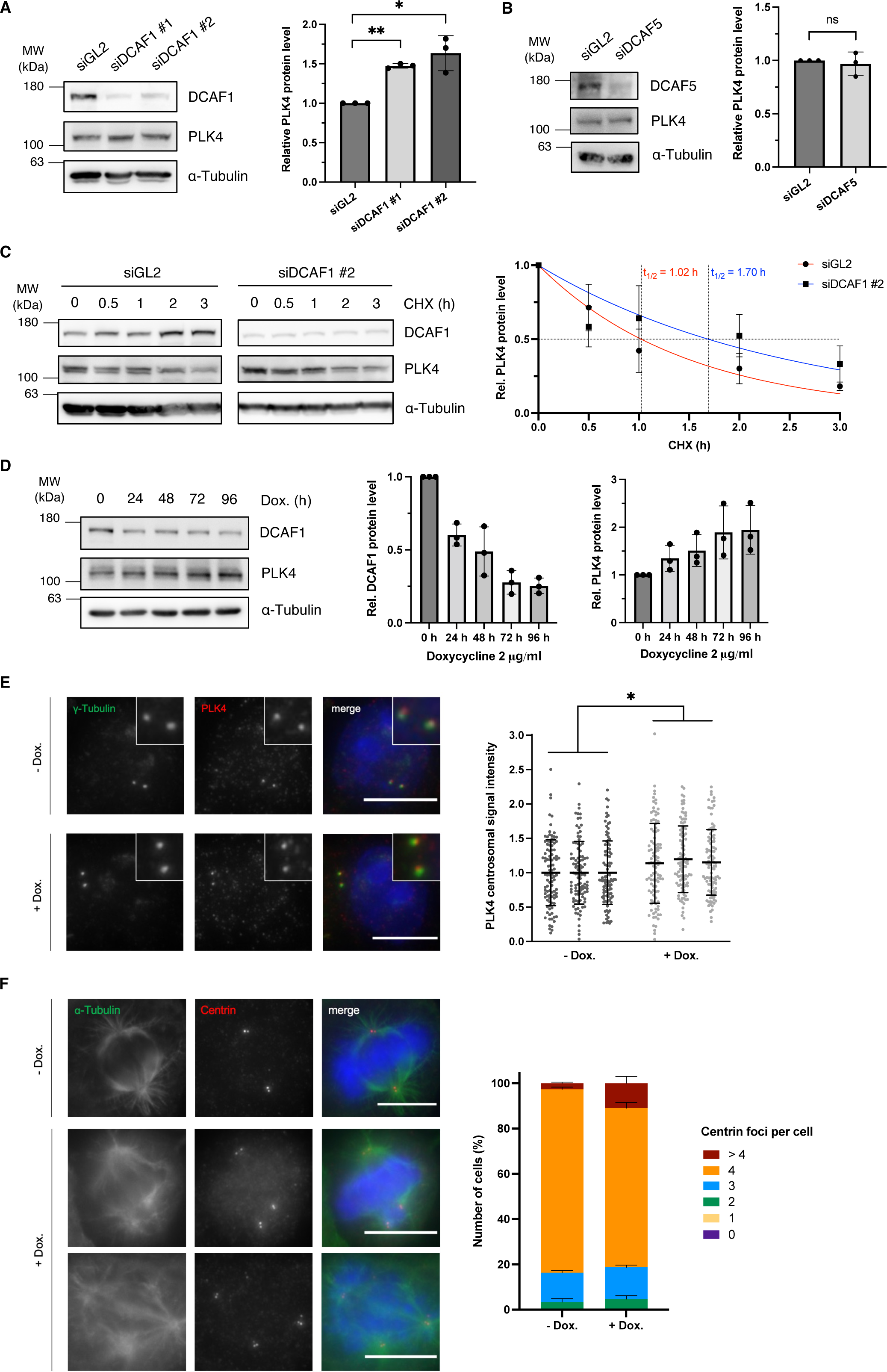
DCAF1 knockdown increases PLK4 protein levels and promotes the formation of supernumerary centrioles in mitosis. **(A)** U2OS cells were transfected twice with siRNA against either GL2 (control) or DCAF1 and harvested 72 h after the first transfection. Protein levels were determined by Western blot analysis. Quantification of relative PLK4/ α-Tubulin signal normalized to siGL2. * p < 0.05, ** p < 0.01, unpaired Student’s t test with Welch’s correction, N = 3. **(B)** HEK293T cells were transfected twice with siRNA against either GL2 (control) or DCAF5 and harvested 72 h after the first transfection. Protein levels were determined by Western blot analysis. Quantification of relative PLK4/ α-Tubulin signal normalized to siGL2. ns p > 0.05, unpaired Student’s t test with Welch’s correction, N = 3. **(C)** U2OS cells were transfected twice with siRNA against either GL2 (control) or DCAF1 and protein synthesis was blocked 72 h after the first transfection by treatment with 100 μg/ml cycloheximide (CHX) for the indicated durations prior to harvest. Protein half-lives were determined by nonlinear fit to a one-phase decay model. N = 3. **(D)** HeLa tet-on shDCAF1 cells were treated with 2 μg/ml doxycycline for the indicated durations prior to harvest. Protein levels were determined by Western blot analysis. N = 3 independent experiments. **(E)** For knockdown of DCAF1, HeLa tet-on shDCAF1 cells were treated with 2 μg/ml doxycycline for 72 h prior to fixation. For immunofluorescence analysis, cells were stained with antibodies against γ-Tubulin and PLK4. Scale bar: 10 μm. Centrosomal signal intensities were quantified and normalized to the untreated control. * p < 0.05, unpaired Student’s t test with Welch’s correction, N = 3 independent experiments, n = 100 centrosomes analyzed per condition. **(F)** For knockdown of DCAF1, HeLa tet-on shDCAF1 cells were treated with 2 μg/ml doxycycline for 72 h prior to fixation. For immunofluorescence analysis, cells were stained with antibodies against α-Tubulin and centrin. Scale bar: 10 μm. The number of centrioles per mitotic cell was determined based on centrin staining. N = 3 independent experiments with n = 100 mitotic cells per condition.

### 2. DCAF1 interacts with and ubiquitylates PLK4

To identify the minimal domain of DCAF1 that binds to PLK4, we used truncated, Flag-tagged fragments of DCAF1 (Cassiday et al., 2015) (Fig. S3B) and performed co-immunoprecipitation experiments. Interestingly, we found that the unstructured acidic domain of DCAF1, located downstream of the WD40 domain at the C-terminal end, mediates the binding to PLK4 (Fig. S3A and B). In addition, DCAF1 strongly binds to a WD40-Acidic motif but not to the WD40 domain alone, suggesting that both domains together might contribute to PLK4 binding.

Next, we aimed at investigating the structure of the DCAF1/PLK4 complex more closely and constructed a model of the complex using alphafold2.0 (Jumper et al., 2021). Recently, Thomä and co-workers determined the structure of CRL4^DCAF1^ by cryo-EM (Mohamed et al., 2021). While the 1.5 MDa structure provides atomic resolution insight into the mechanism of ligase assembly and activation, the DCAF1 C-terminal acidic tail was not observed. Usually, cryo-EM cannot detect unstructured regions, which is consistent with our alphafold2.0 model, suggesting that the acidic tail alone is intrinsically unstructured (Fig. S4). Alphafold2.0 enabled us to construct a model of the DCAF1/PLK4 complex which we compared to the CEP192/PLK4 complex (Fig. 3A). CEP192 is a pericentriolar protein required for nucleation of centrosomal microtubules during mitosis, that strongly binds to the PLK4 PB1-2 (Gomez-Ferreria et al., 2007; Kim et al., 2013; Sonnen et al., 2013). An overlay between the two structures revealed that both CEP192 and DCAF1 bind to the same groove within the PLK4 PB1-2 (Fig. 3B). The structure suggests that a dimer of PLK4 binds to a single chain of the unstructured DCAF1 acidic domain. Binding induces helices formation of the unstructured acidic region: the first helix (D1420-E1436) positioned in a basic groove formed between the PB1 and PB2 of one protomer and the second helix (D1458-E1466) bound to the same basic groove of the second protomer in the PLK4 dimer (Fig. 3C). The model demonstrated that an extended unstructured region (E1467-E1507), downstream of the second helix, wriggles back onto the PB1 surface of the second protomer. A connector region between the two DCAF1 helices forms interactions with a groove mainly of the second protomer. Residues at the PB binding grooves are highly conserved, indicating their importance for protein binding (Fig. 3B). The grooves present positive charge surfaces as visualized by the calculated adaptive Poisson-Boltzmann solver (Fig. 3C).

**Figure 3.**
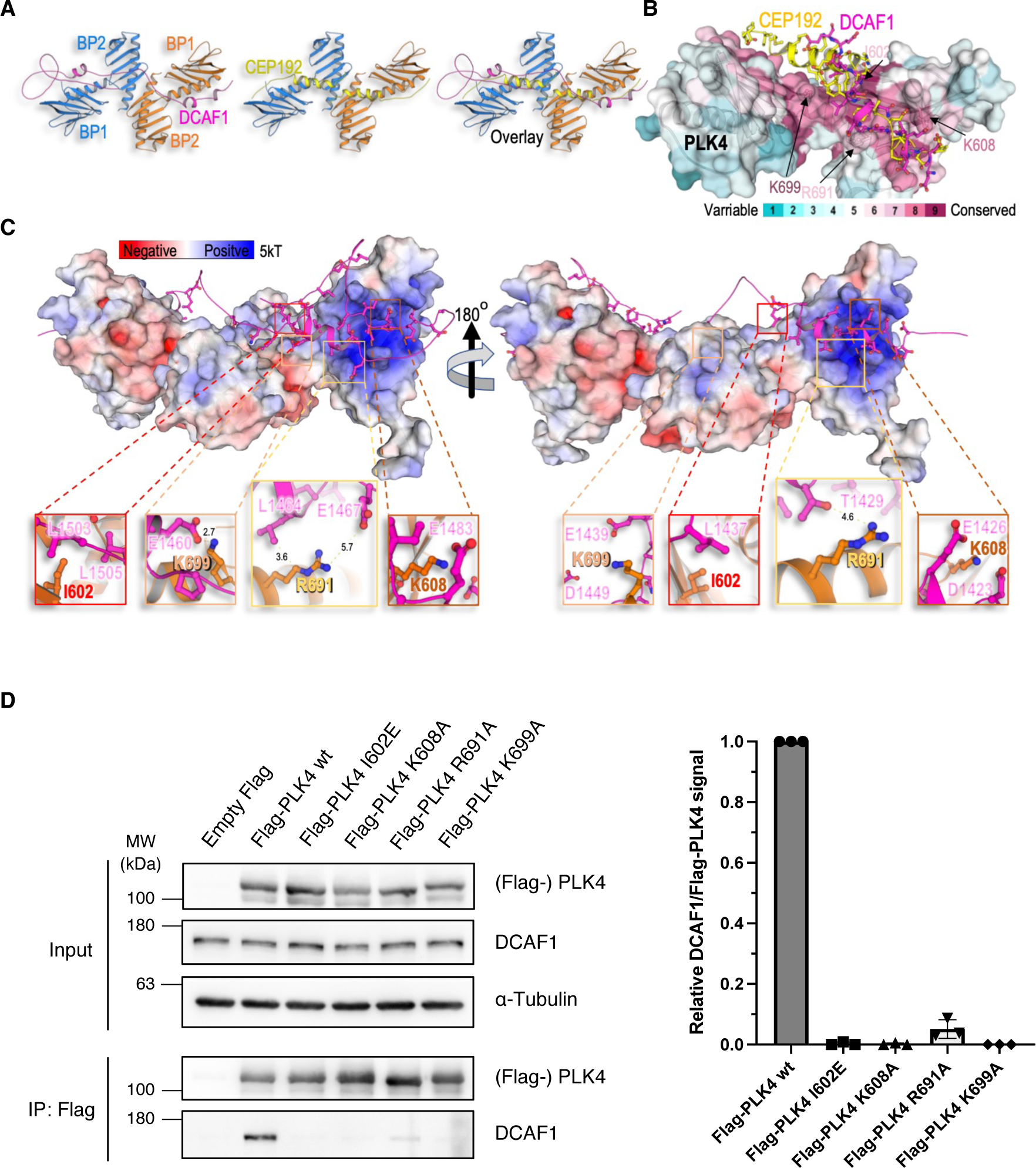
Structure of the DCAF1/PLK4 complex. **(A)** The model of the DCAF1/PLK4 complex was constructed by alphafold2.0. The model was further minimized by Refmac5 (idealization procedure) and examined by structural based mutagenesis and binding assays. Structures of the CEP192/PLK4 complex (PDB 4N7Z), the DCAF1/PLK4 complex and overlay of the two are shown. **(B)** DCAF1 and CEP192 bind a conserved groove in PLK4. Conservation analysis was used to render the conservation level of the residues in the PB1/2 domain of PLK4 which is shown as transparent surface to allow the view of residues that were mutated. CEP192 (yellow) and DCAF1 (magenta) are shown as cartoon with ball-n-sticks residues. **(C)** The surface electrostatic potential was calculated by the Advanced Poisson-Boltzmann Solver (APBS) with the indicated kT. The acidic domain of DCAF1 (magenta cartoon) binds the positive (basic) groove of the PLK4 PB1/2 homodimers. **(D)** Indicated Flag-PLK4 mutants were overexpressed in HEK293T cells for 48 h. Co-precipitated DCAF1 was detected by IP against the Flag tag and subsequent Western blot analysis. Quantification of relative DCAF1/ Flag-PLK4 signal normalized to Flag-PLK4 wt. N = 3.

To confirm the importance of the residues located in the binding grooves for the PLK4-DCAF1 interaction experimentally *in vivo*, PLK4 constructs harbouring point mutations in the PB1 and PB2 (mutations are depicted in bold in Fig. 3C) were generated and transfected into HEK293T cells followed by immunoprecipitation. In alignment with the structural model, we found that binding of DCAF1 to these PLK4 mutants but not to PLK4 wild-type was abolished, suggesting that these amino acids are critical for binding (Fig. 3D).

To assess if the DCAF1-PLK4 interaction is direct or is mediated by another component, we performed ubiquitylation assays in a heterologic environment that lacks such possible mediator component. We used two different *in vitro* ubiquitylation assays: (1) We reconstituted the DCAF1-dependent PLK4 ubiquitylation cascade in *E. coli* (Keren-Kaplan et al., 2012). To circumvent the complexity and the tight regulations of the mammalian system, in the *E. coli*-based, constructed system the E2 enzyme was fused to the substrate receptor DCAF1 and co-expressed with GFP-PLK4, E1 and His_6_-Ubiquitin. We found that PLK4 underwent ubiquitylation when a full cascade was reconstructed. However, strains that expressed only E1, E2-DCAF1 or only PLK4 or a complete cascade but containing a catalytic mutation in the E2 sequence (C86A), did not yield PLK4 ubiquitylation (Fig. 4A). (2) We performed an *in vitro* ubiquitylation assay with recombinant PLK4 and observed ubiquitylation of PLK4 in the presence of UBA1 (E1), UBCH5C (E2) (Han et al., 2020), DCAF1 wild-type but not DCAF1 ΔWD40-ΔAcidic which lacks the PLK4 binding domain (Fig. 4B).

**Figure 4.**
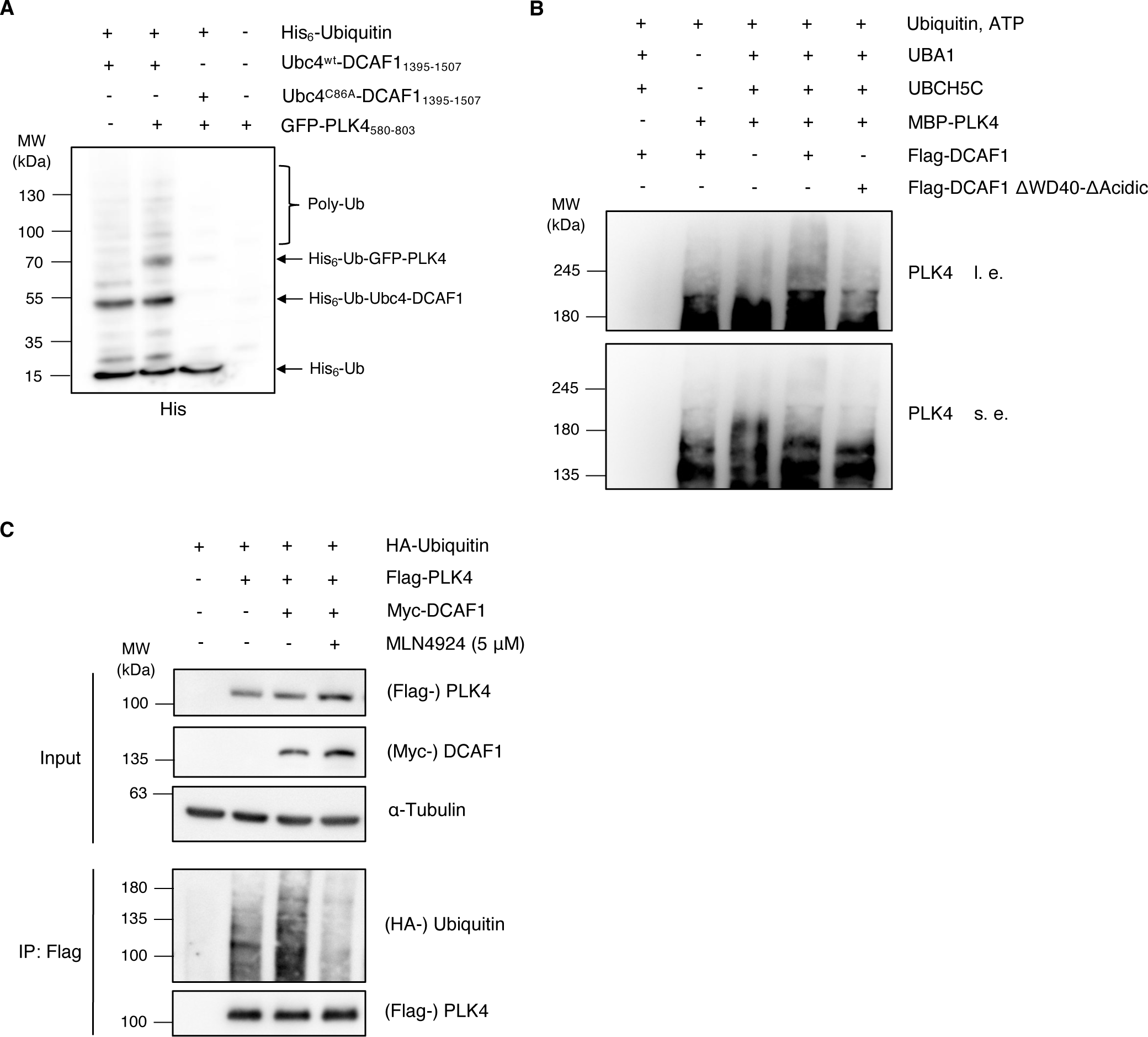
DCAF1 ubiquitylates PLK4 *in vitro* and *in vivo.* **(A)** His-ubiquitin was co-expressed in E. coli together with a GFP-PLK4 construct containing the PB1-PB2 domain of PLK4 and a fusion construct consisting of the DCAF1 acidic domain and the E2 enzyme Ubc4, with or without mutation of the catalytic cysteine. Bacteria cells were harvested and cell lysates were incubated with NTA beads. Ubiquitylated proteins were detected by Western blot. **(B)** Flag-DCAF1/ Myc-CUL4 complexes were expressed in HEK293T cells for 48 h and immobilized on α-Flag M2 beads. In vitro ubiquitylation assays were performed with 200 nm MBP-PLK4, 170 nM UBA1, 1 μM UBCH5C, 30 μM Ubiquitin, 5 mM ATP and immobilized Flag-DCAF1 complexes for 90 min at 37 °C. **(C)** HA-Ubiquitin, Flag-PLK4 and Myc-DCAF1 were overexpressed in HEK293T cells for 24 h with or without inhibition of Cullin-RING E3 ligases by treatment with 5 μM MLN4924 for 5 h prior to harvest. The 26S proteasome was blocked by 10 μM MG132 for 5 h prior to harvest. Flag-PLK4 was immunoprecipitated from cell lysates in the presence of 10 mM *N*-ethylmaleimide using α-Flag M2 beads.

To further corroborate that the regulation of PLK4 by DCAF1 is mediated by ubiquitylation, we also performed *in vivo* ubiquitylation assays. We found that overexpression of DCAF1 led to an increase in PLK4 polyubiquitylation (Fig. 4C). This effect was reversed by inhibition of Cullin-RING E3 ubiquitin ligases with the small-molecule neddylation inhibitor MLN4924. Together, our data show that PLK4 and DCAF1 form complexes *in vivo* and *in vitro*, revealing that PLK4 is a new substrate of the CRL4^DCAF1^ E3 ubiquitin ligase complex.

### 3. DCAF1 binds and ubiquitylates PLK4 predominantly in G2 phase

In mammalian cells, PLK4 is binding to its centriole receptors CEP152 and CEP192, which encircle the proximal end of the parent centriole to initiate centriole duplication at the G1/S phase transition (Cizmecioglu et al., 2010; Hatch et al., 2010; Sonnen et al., 2013; Kim et al., 2013; Park et al., 2014). Activation of PLK4 at the centriole occurs through trans autophosphorylation (Holland et al., 2010). Phosphodegrons generated in response to PLK4 autophosphorylation are recognized by SCF^β−TrCP^ (Guderian et al., 2010; Cunha-Ferreira et al., 2013), which triggers PLK4 degradation (Klebba et al., 2013). As the regulation of PLK4 by CRL4^DCAF1^ is independent of phosphorylation and DCAF1 binds to the PB1-PB2 domain of PLK4, we asked at what time in the cell cycle PLK4 protein levels are regulated by CRL4^DCAF1^ and whether this timing would be different from the regulation of PLK4 by SCF^β-TrCP^ at the G1/S phase transition. To address this question, we expressed PLK4 in cells that were subsequently synchronized and found that DCAF1 strongly interacted with PLK4 during interphase but only weakly in mitosis (Fig. 5A). Similar results were obtained in cells when endogenous DCAF1 was immunoprecipitated instead of PLK4 (Fig. S5). Next, we analyzed whether the cell cycle-regulated interaction between PLK4 and DCAF1 correlates with ubiquitylation of PLK4 by CRL4^DCAF1^. As shown in Fig. 5B using synchronized cells, PLK4 is predominantly ubiquitylated by CRL4^DCAF1^ in G2 phase of the cell cycle. Together, these results suggest that the CRL4^DCAF1^ complex regulates protein levels of PLK4 in G2 phase at a time when SCF^β-TrCP^ E3 ligase activity is low, as shown by Paul et al., 2022.

**Figure 5.**
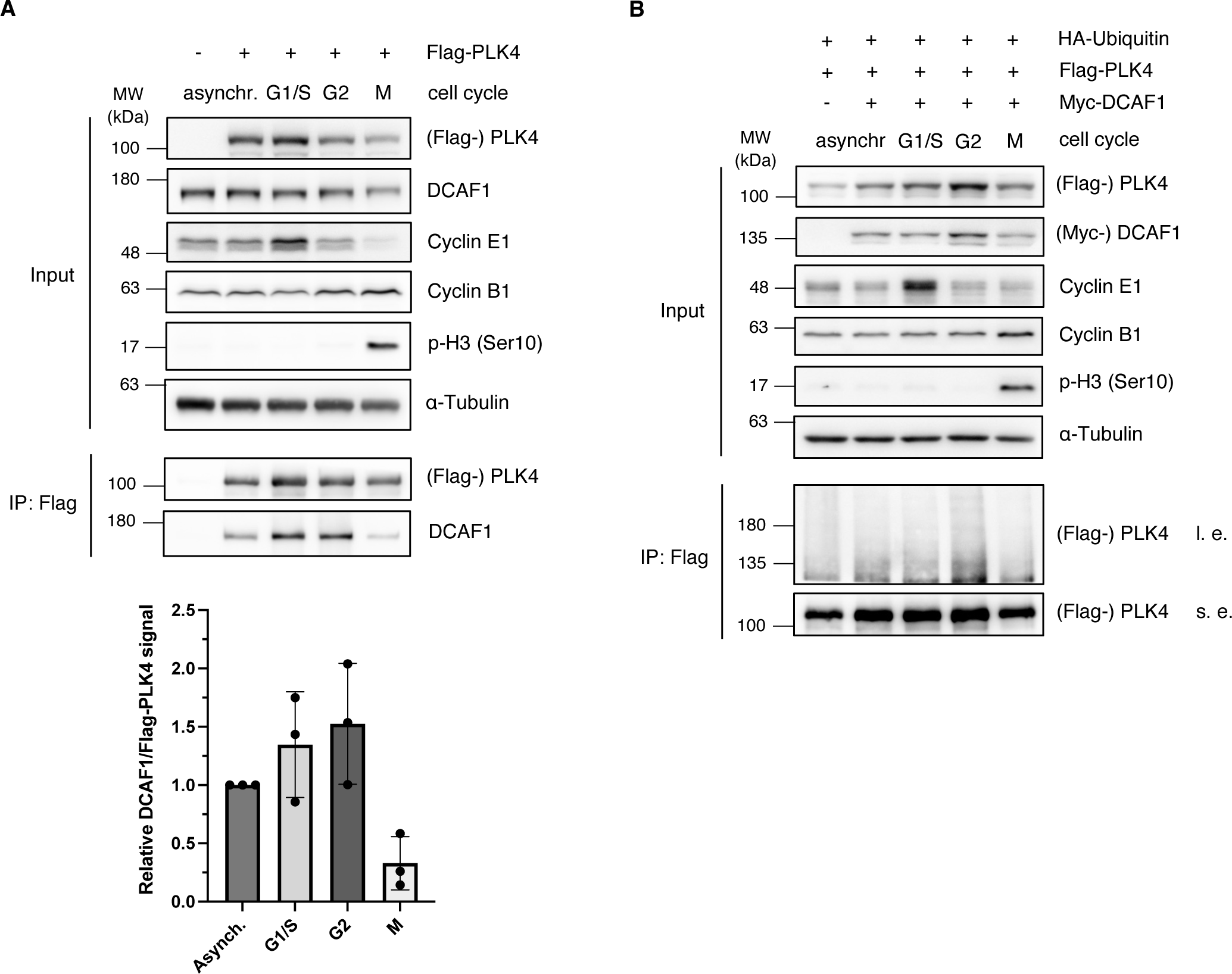
DCAF1 interacts with and ubiquitylates PLK4 predominantly in G2 phase of the cell cycle. **(A)** Flag-PLK4 was overexpressed in HEK293T cells for 48 h. Cells were synchronized in G1/S phase by double thymidine arrest, in G2 phase by CDK1 inhibition with RO-3306 or in M phase by single thymidine and nocodazole arrest, as indicated. Flag-PLK4 was immunoprecipitated from cell lysates using α-Flag M2 beads. Quantification of relative DCAF1/ Flag-PLK4 signal normalized to asynchronous cells. N = 3 independent experiments. **(B)** HA-Ubiquitin, Flag-PLK4 and Myc-DCAF1 were overexpressed in HEK293T cells for 24 h. Cells were synchronized as in (A) and treated with 10 μM MG132 for 5 h prior to harvest. Flag-PLK4 was immunoprecipitated from cell lysates in the presence of 10 mM *N*-ethylmaleimide using α-Flag M2 beads.

### 4. CRL4^DCAF1^ prevents premature centriole disengagement in G2 phase

We then aimed at deciphering a possible function of DCAF1 in regulating PLK4 protein levels. Since we found that DCAF1 ubiquitylates PLK4 in G2 phase, we asked whether it would exert a role in G2 phase to possibly regulate PLK4 functions in mitosis/early G1 phase. During mitosis, the mitotic kinase CDK1-CyclinB binds STIL and prevents formation of the PLK4-STIL complex and STIL phosphorylation by PLK4, thus inhibiting untimely onset of centriole biogenesis (Zitouni et al., 2016). It is conceivable that DCAF1 prevents premature binding of PLK4 to STIL in G2 phase by keeping PLK4 levels low. To find out whether depleting DCAF1 would affect complex formation of PLK4 and STIL, we analyzed the amount of STIL binding to PLK4 in the presence and absence of DCAF1. We found that upon depletion of DCAF1, a higher amount of STIL binds to PLK4 (Fig. 6A). To confirm this finding, instead of depleting DCAF1, an increasing amount of DCAF1 was co-expressed along with PLK4 in cells and the level of STIL binding to PLK4 was assessed. We found that an increase in DCAF1 levels leads to a decrease in the amount of PLK4 binding to STIL (Fig. 6B). These findings indicate that DCAF1 not only has a vital regulatory function for PLK4 itself but also for the interaction between PLK4 and its substrate STIL further downstream. Moreover, depletion of DCAF1 also affects the levels of the PLK4 substrate NEDD1 at the centrosome (Chi et al., 2021) (Fig. S6). After exit from mitosis and entry into G1, the centrioles are “licensed” for a subsequent round of centriole duplication as the engaged centriole pairs lose their tight orthogonal configuration leading to centriole disengagement (Tsou and Stearns, 2006). A premature onset of centriole duplication in the absence of DCAF1 should also result in increased numbers of already disengaged centrioles in G2 phase. Indeed, we found that doxycycline-induced knockdown of DCAF1 leads to a significant higher number of disengaged centrioles in cells synchronized in G2 phase (Fig. 6C) and a decrease in the levels of the centriole-linker protein rootletin (Fig. 6D), indicating a precocious centriole disengagement. These results suggest that controlled PLK4 protein levels in G2 phase are necessary to prevent unscheduled centriole duplication.

**Figure 6.**
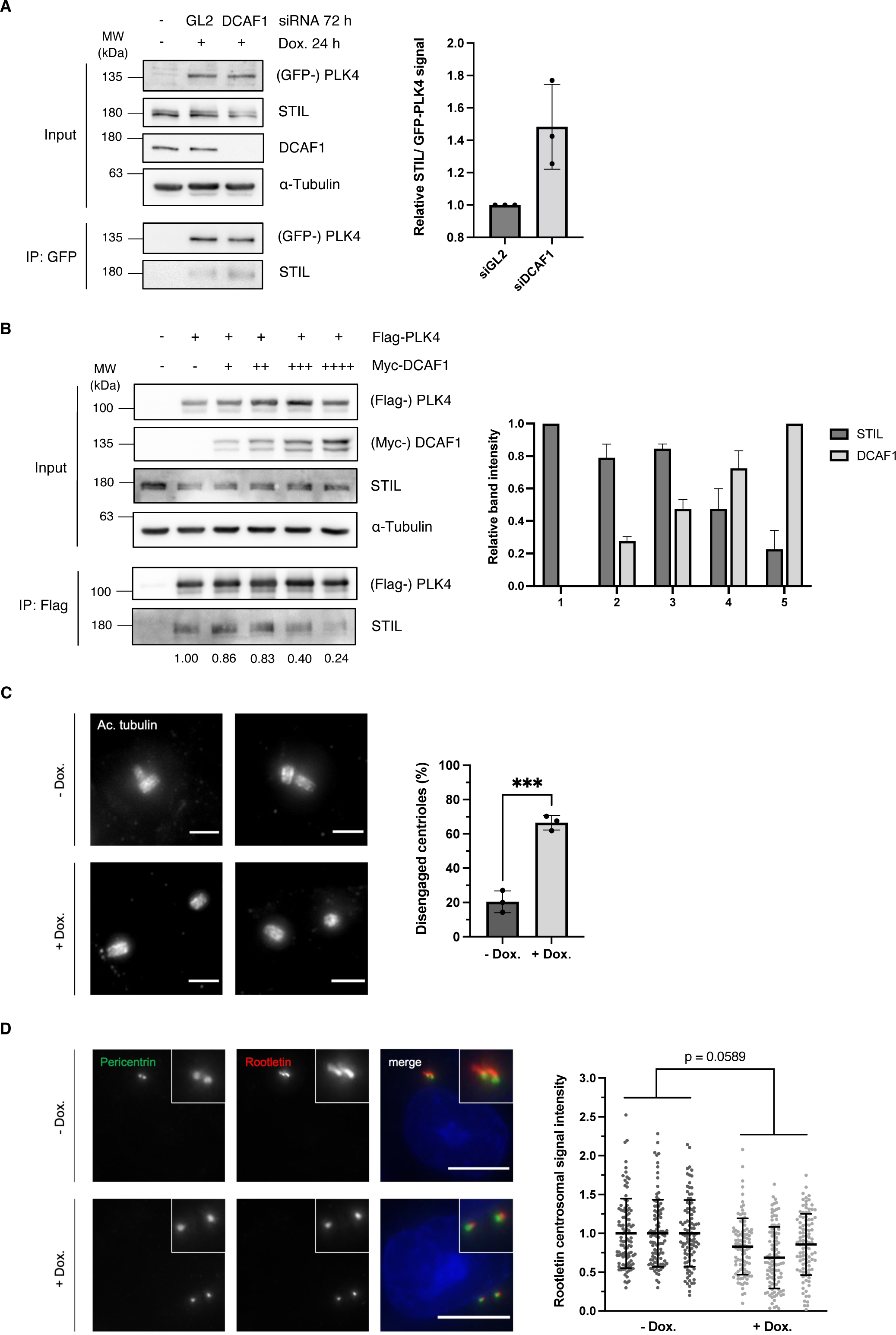
DCAF1 regulates the interaction of PLK4 with its substrate STIL and is required to prevent premature centriole disengagement in G2 phase. **(A)** HeLa tet-on GFP-PLK4 cells were transfected twice every 24 h with 40 nM siRNA targeting GL2 (control) or DCAF1. Overexpression of GFP-PLK4 was induced by treatment with 2 μg/ml doxycycline 24 h prior to harvest. GFP-PLK4 was immunoprecipitated from cell lysates using GFP-trap beads. Quantification of relative STIL/ GFP-PLK4 signal normalized to GL2 (control). N = 3 independent experiments. **(B)** Flag-PLK4 was overexpressed in HEK293T cells together with different amounts of Myc-DCAF1 for 48 h. Co-precipitated STIL was detected by IP against the Flag tag and subsequent Western blot analysis. Quantification of relative STIL/ Flag-PLK4 and Myc-DCAF1 signal. N = 3 independent experiments. **(C)** For knockdown of DCAF1, HeLa tet-on shDCAF1 cells were treated with 2 μg/ml doxycycline for 72 h prior to fixation. G2 arrest was induced by treatment with the CDK1 inhibitor RO-3306 for 18 h prior to fixation. Representative expansion microscopy images of centrioles stained against acetylated tubulin. Scale bar: 3 μm (physical scale), 0.68 μm (biological scale). Quantification of percentage of cells with disengaged centrioles in G2 phase. N = 3 independent experiments with n = 37, 42 and 40 cells analyzed per condition. Unpaired Student’s t test with Welch’s correction. **(D)** For knockdown of DCAF1, HeLa tet-on shDCAF1 cells were treated with 2 μg/ml doxycycline for 72 h prior to fixation. G2 arrest was induced by treatment with the CDK1 inhibitor RO-3306 for 18 h prior to fixation. For immunofluorescence analysis, cells were stained with antibodies against pericentrin and rootletin. Scale bar: 10 μm. Centrosomal signal intensities were quantified and normalized to the untreated control. Unpaired Student’s t test with Welch’s correction, N = 3 independent experiments, n = 100 centrosomes analyzed per condition.

## Discussion

Extensive research has shown that centrosome number control is critical for the maintenance of genomic integrity. In this study, we introduce an additional layer of regulation that allows timely removal of PLK4, the master regulator of centriole duplication (Nigg and Holland, 2018; Goundiam and Basto, 2021). Our data indicate that at least two pathways exist that regulate PLK4 protein levels and restrict centriole duplication to once per cell cycle, in order to prevent excess centrosome number. These pathways are governed by two distinct ubiquitylating enzymes, one where the SCF^β-TrCP^ E3 ubiquitin ligase directly recognizes phosphorylated S285/T289 on PLK4 (Guderian et al., 2010; Holland et al., 2010; Cunha-Ferreira et al., 2013; Klebba et al., 2013) and one which is independent of PLK4 autophosphorylation and mediated by the CRL4^DCAF1^ E3 ubiquitin ligase complex. Interestingly, the two pathways do not only differ in respect to the phosphorylation status of PLK4 but also in the different phases of the cell cycle where they regulate PLK4 protein levels. Whereas SCF^β-TrCP^ controls PLK4 levels at the G1/S phase transition, we find that CRL4^DCAF1^ targets PLK4 during G2 phase to control its levels during a different phase of the cell cycle. Recently, Paul et al. have shown by using a fluorescent biosensor to quantitatively measure β-TrCP activity, that β-TrCP is highly active during quiescent G0 state, moderately active in G1 phase and the least active during S and G2 phase (Paul et al., 2022). Our own data reveal that DCAF1 binds PLK4 predominantly in G1/S and G2 phase (Fig. 5A) but this binding leads to a stronger ubiquitylation of PLK4 only in G2 phase (Fig. 5B). From procentriole assembly throughout S phase until late mitosis, the procentrioles remain in a tight, near orthogonal association with their parental centrioles. This connection is lost in late mitosis/ early G1, during centriole disengagement. The centriole-procentriole engagement is thought to prevent unscheduled procentriole assembly. We propose that PLK4 protein levels have to be controlled especially in G2 phase by CRL4^DCAF1^-mediated ubiquitylation and degradation to prevent premature centriole disengagement in G2, a process that is critical for licensing the subsequent round of centrosome duplication (Tsou and Stearns, 2006). Untimely onset of centriole duplication is prevented by the mitotic kinase CDK1-CyclinB that binds STIL and prevents formation of the PLK4-STIL complex and STIL phosphorylation by PLK4 (Zitouni et al., 2016). Our findings imply that CRL4^DCAF1^ binds and ubiquitylates PLK4 in G2 phase but not in mitosis. As ubiquitylation of PLK4 by CRL4^DCAF1^ causes degradation of PLK4, we propose a mechanism where low PLK4 levels prevent binding and activation of the PLK4 substrate STIL, thus impeding premature initiation of centriole duplication. CDK1-CyclinB may prevent STIL-PLK4 interaction by binding STIL in a kinase-independent fashion through the same region as PLK4 (Zitouni et al., 2016). It is conceivable that increasing amounts of PLK4 during mitosis, when the interaction between CRL4^DCAF1^ and PLK4 is weak, might capture STIL from binding to CDK1, initiating the interaction between PLK4 and STIL. Thus, the regulation of PLK4 protein levels by CRL4^DCAF1^ represents another important pathway to control PLK4 activation and binding to STIL.

We found that the interaction between DCAF1 and PLK4 has also interesting structural aspects. PLK4 is a structurally divergent PLK family member characterized by a single polo box (PB3), which is capable of intermolecular homodimerization and a conserved central region called ‘cryptic polo box’ (PB 1-2), which is necessary for its functions in centriole duplication. Since we found that DCAF1 binds PB1-2 of PLK4 (Fig. S2), we compared our model of the DCAF1/PLK4 complex structure (Fig. 3A and B) with the CEP192/PLK4 and CEP152/PLK4 complexes, since both CEP192 and CEP152 were previously shown to also bind PB1 and PB2 of PLK4 (Sonnen et al., 2013; Kim et al., 2013). Acidic helical regions of CEP152 and CEP192 bind to PB1-2 of PLK4 in opposite directions (Park et al., 2014). Interestingly, we found that PLK4 binds CEP192 and DCAF1 in a very similar manner via a conserved groove within the PLK4 PB1-PB2 domain (Fig. 3B). However, while each of CEP152 and CEP192 present a short acidic helix, DCAF1 has two short acidic helices connected by a flexible linker that allows them both to bind simultaneously. The first region (D1420-E1436) binds with the same orientation as CEP192, whereas the second region (D1458-E1466) interacts with the same orientation as CEP152. It would be interesting to determine whether DCAF1 and CEP152/CEP192 bind mutually exclusive to PLK4 or whether they collaborate with PLK4 for the regulation of centriole duplication.

A common structural mechanism for protein-protein interactions is often achieved by β-sheet assembly, where one interaction partner donates a single β-strand and the other partner donates a β-sheet that lacks one or few strands. Such phenomenon was previously described as the structural basis for the binding of the Merlin FERM domain to DCAF1 (Li et al., 2014). The terminal β-strand of DCAF1 (amino acids DIILSLN in the β-B) forms a β-sheet with βF5 of the FERM domain. Careful examination of the alphafold2.0 model of the DCAF1/PLK4 complex suggests that a similar interaction takes place for the same DCAF1 sequence to extend a PLK4 β-sheet of PB1.

Targeted protein degradation represents an emerging therapeutic modality with the potential to tackle disease-causing proteins that have in the past been highly challenging to target with conventional small molecules. Our results pave the way for the development of CRL4^DCAF1^-dependent PROTACs or molecular glue degraders (Bekes et al., 2022; Levin-Kravets et al., 2021) to target PLK4 degradation as a novel modality for cancer therapy.

## Materials and Methods

### Cell lines, cell culture and cell cycle synchronization

HEK293T (catalog no.: ACC 635, DSMZ Braunschweig, Germany), HeLa (ATCC CCL-2), HeLa tet-on shDCAF1 and U2OS cells (ATCC HTB-96) were cultured in Dulbeccós modified Eaglés medium containing 4.5 g/l glucose (Gibco, catalog no.: 41965-039) supplemented with 10 % fetal bovine serum (Gibco, catalog no.: 10270-106) and 1 % penicillin/ streptomycin (Sigma, catalog no.: P0781) at 37 °C and 5 % CO_2_.

For generation of a stable cell line for conditional DCAF1 knockdown, HeLa S/A cells (O. Gruss, ZMBH Heidelberg, Germany) were transiently co-transfected with a pBi-9 vector containing the siDCAF1 shRNA sequence and the pCAGGS-flpE vector (O. Gruss, ZMBH Heidelberg, Germany). Antibiotic selection for successful integration of the construct was performed using 2.5 μg/ml puromycin (Sigma-Aldrich) and 10 μM ganciclovir (Alpha Diagnostic Intl. Inc.). DCAF1 knockdown was induced by addition of 2 μg/ml doxycycline (Sigma-Aldrich, catalog no.: D9891) for 72 h.

Cullin-RING E3 ubiquitin ligases were inhibited by addition of 5 μM MLN4924 (Cell Signaling Technology, catalog no.: 85923) and the 26S proteasome was blocked with 10 μM MG132 (Sigma-Aldrich, catalog no.: C2211) for 5 h prior to harvest.

For double thymidine arrest of HEK293T cells in G1/S phase, cells were treated with 2 mM thymidine (Santa Cruz Biotechnology, catalog no.: sc-296542A) for 18 h, released for 9 h and arrested again for 16 h. For single thymidine and nocodazole arrest in mitosis, cells were treated with 2 mM thymidine for 20 h, released for 2 h and treated with 100 ng/ml nocodazole (Merck) for 16 - 17 h. For G2 phase arrest of HeLa or HEK293T cells, cells were treated with 10 μM RO-3306 for 18 h.

### Plasmids, cloning and mutagenesis

pCMV-3Tag1A-Flag-PLK4 full length and fragments have been described previously (Cizmecioglu et al., 2010). pCMV-3Tag1A-Flag-PLK4 mutants were generated by site-directed mutagenesis. pCMV-Sport6-Flag-DCAF1 full length and fragments were a gift from Prof. Vicente Planelles (Cassiday et al., 2015). pCMV-3Tag2C-Myc-DCAF1 was generated by subcloning using EcoRI and XhoI. For the generation of a stable cell line for inducible knockdown of DCAF1, shRNA constructs were cloned to the pBI9 vector (O. Gruss, ZMBH Heidelberg, Germany) using BsaI. Constructs for expression and detection of ubiquitylated PLK4 in *E. coli* were generated by Gibson assembly (Gibson et al., 2009). EGFP-PLK4_580-808_ was subcloned into pET22b and the His_6_-tag was removed. DCAF1_1395-1507_ was subcloned into pGEN13 upstream of and in-frame with Ubc4 (Keren-Kaplan et al., 2012). The C86A mutation at the catalytic cysteine of Ubc4 was also generated by Gibson assembly.

### Plasmid and siRNA transfections

HEK293T cells were transfected with plasmid DNA using polyethylenimine (Polysciences) at a final concentration of 5 μg/ml for 24 h or 48 h. HeLa and HEK293T cells were transfected with siRNA using Lipofectamine 2000 (Invitrogen). Cells were transfected with 40 nM of siRNA 24 h and 48 h after seeding and further cultivated for another 48 h after the second siRNA transfection.

The following siRNA sequences were used:

GL2 5’-CGUACGCGGAAUACUUCGAdTdT-3’

DCAF1 #1 5’-UCACAGAGUAUCUUAGAGAdTdT-3’ (Nakagawa et al., 2015)

DCAF1 #2 5’-CGGAGUUGGAGGAGGACGAUUdTdT-3’ (Hakata et al., 2014)

DCAF5 5‘-GCAGAAACCUCUACAAGAAdTdT-3’ (Ambion, silencer select)

### Cell lysis, co-immunoprecipitation and Western blot analysis

Cell lysates were prepared as described previously (Hanle-Kreidler et al., 2022). Briefly, cells were harvested and washed with ice-cold PBS. For Western blot analysis, cell pellets were lysed in RIPA buffer (50 mM Tris-HCl pH 7.4, 1 % NP-40, 0.5 % sodium deoxycholate, 0.1 % SDS, 150 mM NaCl, 2 mM EDTA, 50mM NaF) and for immunoprecipitations cell pellets were lysed in NP-40 buffer (40 mM Tris-HCl pH 7.5, 150 mM NaCl, 5 mM EDTA, 10 mM β-glycerophosphate, 5 mM NaF, 0.5 % NP-40). Both buffers were supplemented with 1 mM DTT, 10 μg/ml l-1-tosylamido-2-phenylethyl chloromethyl ketone, 5 μg/ml tosyl lysyl chloromethyl ketone, 0.1 mM Na_3_VO_4_, 1 μg/ml aprotinin, 1 μg/ml leupeptin and 10 μg/ml trypsin inhibitor from soybean. Cell lysates were incubated on ice for 30 min and centrifuged for 20 min at 16,100 g. For SDS-PAGE, cell extracts were mixed with 2x Laemmli buffer and incubated for 5 min at 95 °C. For immunoprecipitations of Flag-tagged proteins, 3 – 6 mg of cell extract and 20 μl of α-Flag M2 affinity bead suspension (Sigma, catalog no.: A2220) were used. The beads were prepared by washing twice with TBS, once with glycine buffer (0.1 M glycine-HCl pH 3.5) and three times with TBS. Cell extracts were incubated with beads for 3 h or overnight on a rotating wheel at 4 °C. Prior to elution, the beads were washed 3 times with NP-40 buffer. For elution, the beads were incubated with 3x Flag peptide (Thermo Fisher Scientific, catalog no.: A36805) for 30 min on ice with short vortexing every 5 – 10 min. Eluates where mixed with Laemmli buffer and denatured for 5 min at 95 °C. Proteins were resolved by SDS-PAGE and detected by chemiluminescence after Western blot. For immunoprecipitations of GFP-tagged proteins, cell extracts were incubated with GFP-trap beads. For endogenous immunoprecipitations, cell extracts were incubated with 4 μg of mouse anti-DCAF1 (Santa Cruz Biotechnology, catalog no.: sc-376850) antibody overnight on a rotating wheel. Protein G sepharose beads were added for 2h at 4 °C. The beads were washed three times with NP-40 buffer and proteins were eluted by incubation with 2x Laemmli buffer for 5 min at 95 °C. Samples were analyzed by SDS-PAGE and Western blot.

### Cycloheximide (CHX) chase

For the analysis of PLK4 protein stability in the presence or absence of DCAF1, U2OS cells were transfected twice with siRNA against GL2 (control) or DCAF1 using Lipofectamine 2000. 72 h after the first transfection, 100 μg/ml cycloheximide (ChemCruz, Santa Cruz Biotechnology, catalog no.: sc-3508) was added to block protein synthesis. Samples were harvested at different time points and analyzed by SDS-PAGE and Western blot.

### *In vivo* ubiquitylation assay

HEK293T cells were transfected with the indicated plasmids for 24 h. Cells were treated with 10 μM proteasome inhibitor MG132 for 5 h prior to harvest. Cullin-RING E3 ligases were inhibited by treatment with 5 μM MLN4924 for 5 h prior to harvest. Cells were harvested and lysed in NP-40 buffer supplemented with 10 mM *N*-ethylmaleimide (Sigma, catalog no.: E3876). Flag-immunoprecipitations were performed as described previously. Whole cell extracts and eluates where mixed with Laemmli buffer and denatured for 5 min at 95 °C. Proteins were resolved by SDS-PAGE and detected by chemiluminescence after Western blot.

### *In vitro* ubiquitylation assays

For *in vitro* ubiquitylation assays using a reconstituted ubiquitylation system in E. coli, T7 Express competent E. coli were co-transformed with the indicated constructs and grown in 1 l LB cultures. 0.1 M IPTG was added for induction at OD_600_ = 1.8 and the bacteria were further grown at 18 °C for 12 h. Cells were harvested by centrifugation and lysed with lysozyme in the presence of serine protease inhibitor (AEBSF). Following sonication and high-speed centrifugation, the soluble fraction was incubated with Ni-NTA beads and washed three times in batch with 150 mM NaCl and 50 mM Tris-HCl pH 7.5. SDS loading buffer was added and the samples were incubated for 10 min at 70 °C prior to SDS-PAGE and Western blot analysis.

Recombinant MBP-PLK4 (Cizmecioglu et al., 2010) was expressed in and purified from E. coli Rosetta (DE3). UBA1, UBCH5C and Ubiquitin were kind gifts from Frauke Melchior (University of Heidelberg). Flag-DCAF1/ Myc-CUL4 complexes were co-expressed in HEK293T cells for 48 h and immunoprecipitated using α-Flag M2 affinity beads as described previously, but proteins were not eluted after washing. Ubiquitylation reactions were performed in 50 mM Tris-HCl pH 7.5, 100 mM NaCl, 10 mM MgCl_2_, 0.05 % NP-40, 1 mM DTT, 0.1 mM Na_3_VO_4_, 1 μg/ml aprotinin, 1 μg/ml leupeptin and 10 μg/ml trypsin inhibitor from soybean. Flag-DCAF1/ Myc-CUL4 complexes immobilized on α-Flag M2 beads were combined with 200 nM MBP-PLK4, 170 nM UBA1, 1 μM UBCH5C, 30 μM Ubiquitin and 5 mM ATP in 20 μl total volume assay buffer. Samples were incubated for 90 min at 37 °C and 400 rpm. Reactions were terminated by the addition of 2x LDS buffer and incubation for 10 min at 72 °C. Samples were analyzed by SDS-PAGE and Western blot.

### Antibodies

Proteins were detected by Western blot using following antibodies: α-DCAF1 (C-8, catalog no.: sc-376850), α-CUL4 (H-11, catalog no.: sc-377188) and α-cyclin E1 (HE12, catalog no.: sc-247) antibodies were purchased from Santa Cruz Biotechnology; α-tubulin (catalog no.: T5168), α-Flag (catalog no.: F3165) and α-polyhistidine-peroxidase (catalog no.: A7058) antibodies were purchased from Sigma; α-DDB1 (catalog no.: A300-462A) and α-STIL (catalog no.: A302-442A) were purchased from Bethyl; α-phospho-H3 (Ser10) (catalog no.: 06-570) antibody was purchased from Merck; α-HA tag (16B12) antibody was purchased from Babco; α-β-TrCP (D13F10, catalog no.: 4394S) antibody was purchased from Cell Signaling Technology, the α-PLK4 and α-cyclin B1 antibodies have been described previously (Cizmecioglu et al., 2010) (Hoffmann et al., 1993) and the α-DCAF5 antibody was generated by Innovagen AB, Lund, Sweden.

For immunofluorescence, α-pericentrin (catalog no.: ab4448) antibody was purchased from Abcam; α-centrin antibody (catalog no.: 04-1624) was purchased from Merck; α-rootletin (C-2, catalog no.: sc-374056) antibody was purchased from Santa Cruz Biotechnology; α-tubulin (catalog no.: T5168) and γ-tubulin (GTU-88, catalog no.: T6557) antibodies were purchased from Sigma-Aldrich and the α-DCAF1 antibody is mentioned above.

### Immunofluorescence microscopy

Cells grown on coverslips were washed with PBS and fixed with ice-cold methanol for 10 min at - 20 °C. Cells were washed with PBS again and blocked with 3 % BSA and 0.05 % Triton X-100 in PBS (PBS-BT) for 30 min at room temperature. Primary and secondary antibodies were diluted in PBS-BT and the antibody incubations were performed for 1 h at room temperature with three washing steps in between. After washing with PBS another three times, coverslips were mounted onto glass microscope slides using ProLong^TM^ Gold Antifade (Molecular Probes by Life Technologies) with DAPI. Cells were imaged using the Zeiss Observer.Z1 inverted motorized microscope and images were processed using Fiji software.

### Ultrastructure expansion microscopy (U-ExM)

Expansion microscopy was performed according to the previously published protocol by (Gambarotto et al., 2019). Cells on coverslips were fixed using ice-cold methanol for 10 min at – 20 °C. Cells were washed with PBS and incubated in a 0.7 % formaldehyde and 1 % acrylamide solution for 4 h at 37 °C. Gel polymerization on the coverslips was performed using a gelation solution (23 % w/v sodium acrylate, 10 % w/v acrylamide, 0.1 % w/v BIS, 0.5 % TEMED, 0.5 % ammonium persulfate in PBS). Gels were allowed to polymerize for 5 min on ice before coverslips were transferred to 37 °C for 1 h. After complete polymerization, coverslips with gels were incubated in denaturation buffer (200 mM SDS, 200 mM NaCl, 50 mM Tris in ddH_2_O) for 15 min at room temperature to detach the gels from the coverslips. Gels were transferred to 1.5 ml microcentrifuge tubes filled with denaturation buffer and incubated for 90 min at 95 °C. Gels were then transferred to beakers filled with ddH_2_O and washed with ddH_2_O additional two times for 10 min each. Before antibody labeling, gels were shrank by replacing ddH_2_O with PBS, washed twice with PBS for 15 min each and then transferred into a six-well plate. Primary antibodies were diluted in 2 % BSA in PBS and added to the gels. Primary antibody staining was performed in the six-well plate overnight at 4 °C with agitation. Gels in the six-well plate were washed three times for 10 min each with 0.1 % Tween-20 in PBS (PBST) at room temperature with agitation. Secondary antibodies were diluted in 2 % BSA in PBS, added to the gels and incubated for 2.5 h at 37 °C with agitation and protected from light. Gels were washed in the six-well plate as before, transferred to beakers filled with about 1000 ml ddH_2_O and incubated in the dark until mounting and imaging.

### Mass spectrometry analysis of Flag-PLK4 interaction partners

For identification of Flag-PLK4-interacting proteins, Flag-PLK4 elution fractions were resolved by SDS-PAGE and co-precipitating proteins were detected in gel by staining with Colloidal Coomassie. Analysis was performed by M. Schnölzer/ DKFZ Protein Analysis Facility (Heidelberg) as described (Kratz et al., 2015). In brief, the gel lanes were cut into slices, digested with trypsin after reduction and alkylation of cysteines. Tryptic peptides were analyzed by nanoLC-ESI-MS/MS using a nanoAcquity UPLC system (Waters GmbH) coupled online to an LTQ Orbitrap XL mass spectrometer (Thermo Scientific). Data were acquired by scan cycles of one FTMS scan with a resolution of 60000 at m/z 400 and a range from 300 to 2000 m/z in parallel with six MS/MS scans in the ion trap of the most abundant precursor ions. Instrument control, data acquisition and peak integration were performed using the Xcalibur software 2.1 (Thermo Scientific, Bremen, Germany). Database searches were performed against the SwissProt database with taxonomy “human” using the MASCOT search engine (Matrix Science, London, UK; version 2.2.2). MS/MS files from the individual gel slices of each lane were merged into a single search. Peptide mass tolerance for database searches was set to 5 ppm and fragment mass tolerance was set to 0.4 Da. Significance threshold was p<0.01. Carbamidomethylation of cysteine was set as fixed modification. Variable modifications included oxidation of methionine and deamidation of asparagine and glutamine. One missed cleavage site in case of incomplete trypsin hydrolysis was allowed.

### Statistical analysis

All statistical analyses were performed with GraphPad Prism, version 9 (GraphPad Software, Inc). Data were collected from three independent experiments and represented as individual values or as mean ± SD. Statistical significance was analyzed by unpaired, two-tailed, Student’s t test with Welch’s correction as indicated in the figure legend. P values below 0.05 were considered statistically significant (ns p > 0.05, * p < 0.05, ** p < 0.01, *** p < 0.001 and **** p < 0.0001).

## Acknowledgements

We are grateful to Vicente Planelles (University of Utah, USA) and Frauke Melchior (ZMBH, University of Heidelberg, Germany) for providing reagents. Members of our labs are thanked for their comments on the manuscript.

This work was supported by the DKFZ PhD program and the Deutsche José Carreras-Leukämiestiftung (DJCLC R09/30f) both to I.H., grants from the DKFZ-MOST cooperation program (Ca196) to I.H. and G.P. and ICRF 940283 to G.P.

---

**Supplemental Figure 1.**
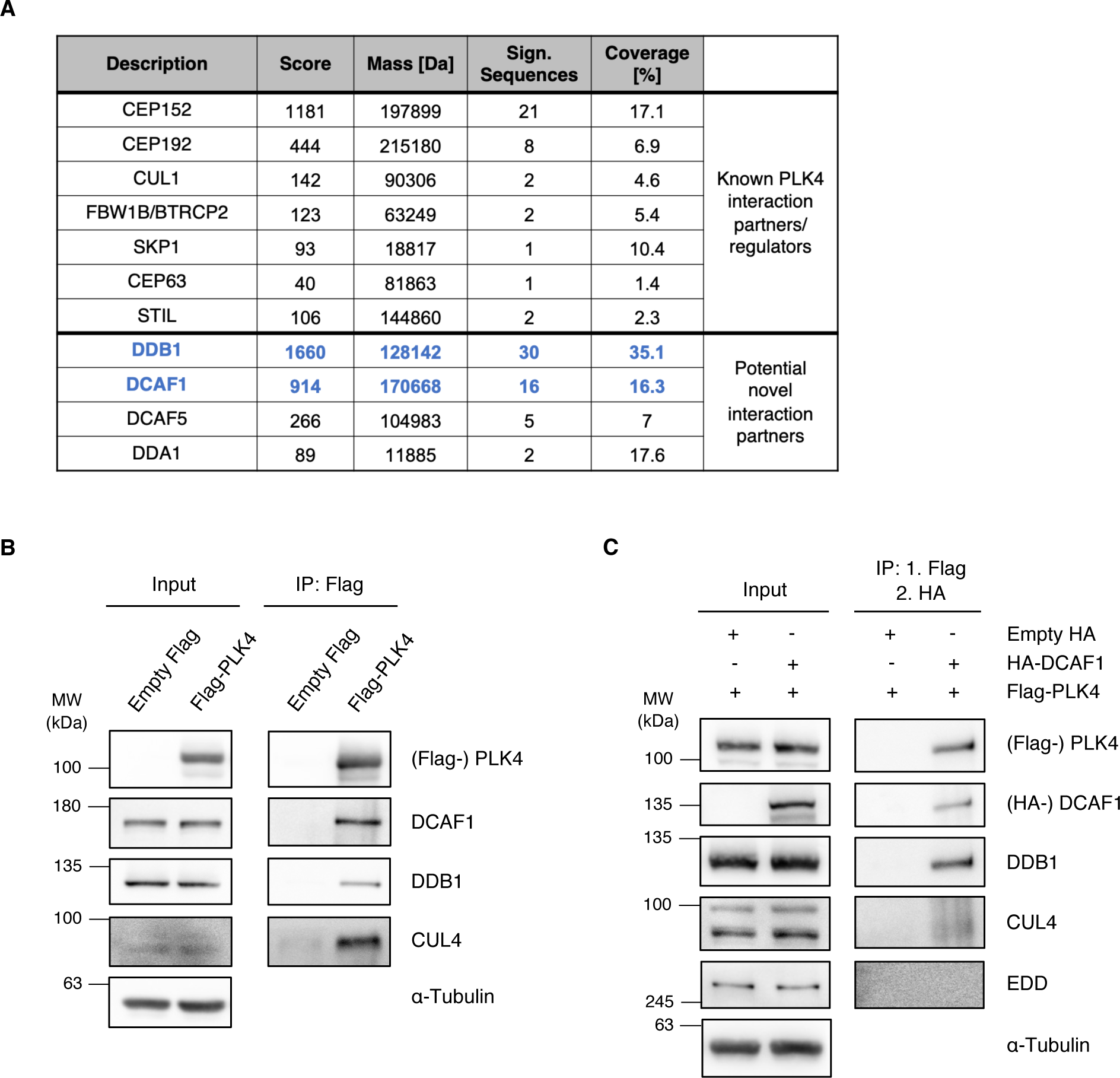
**(A)** Mass spectrometry analysis of Flag-PLK4 IP samples identified known PLK4 interaction partners, substrates or regulators and potential novel interaction partners. MASCOT score, molecular weight (mass), identified protein sequence and coverage of selected candidates are shown. **(B)** Flag-PLK4 was overexpressed in HEK293T cells for 48 h. Co-precipitated E3 ligase complex components DCAF1, DDB1 and CUL4 were detected by IP against the Flag tag and subsequent Western blot analysis. **(C)** Flag-PLK4 and HA-DCAF1 were overexpressed in HEK293T cells for 48 h. Presence of DBB1 and CUL4 in the same complex was analyzed by IP against the Flag tag, followed by Flag peptide elution, subsequent IP against the HA tag using the eluates and Western blot analysis of the double IP samples.

**Supplemental Figure 2.**
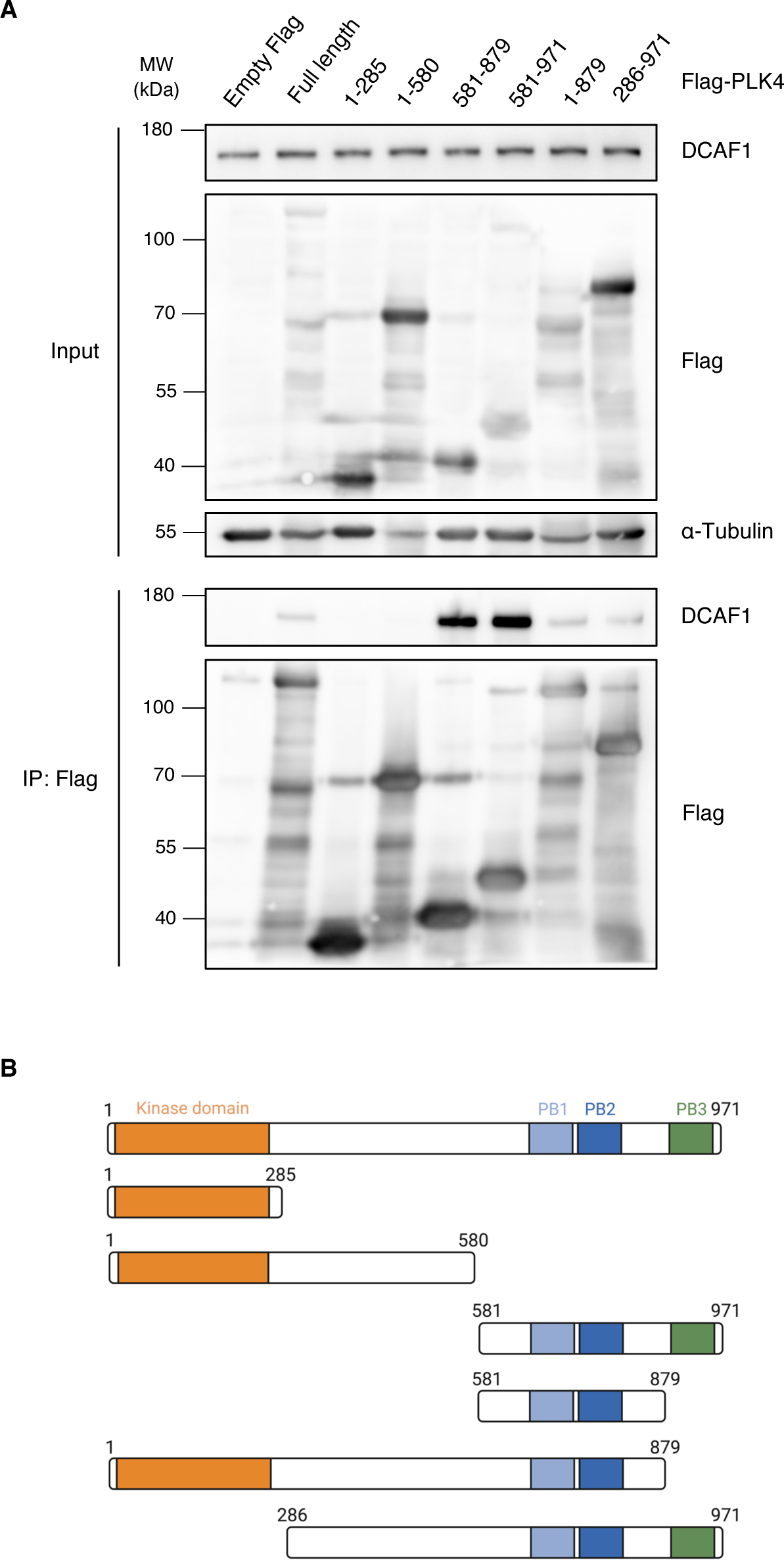
**(A)** Flag-PLK4 full length or different truncated fragments were overexpressed in HEK293T cells for 48 h. Co-precipitated DCAF1 was detected by IP against the Flag tag and subsequent Western blot analysis. **(B)** Overview of the different Flag-PLK4 fragments used in (A).

**Supplemental Figure 3.**
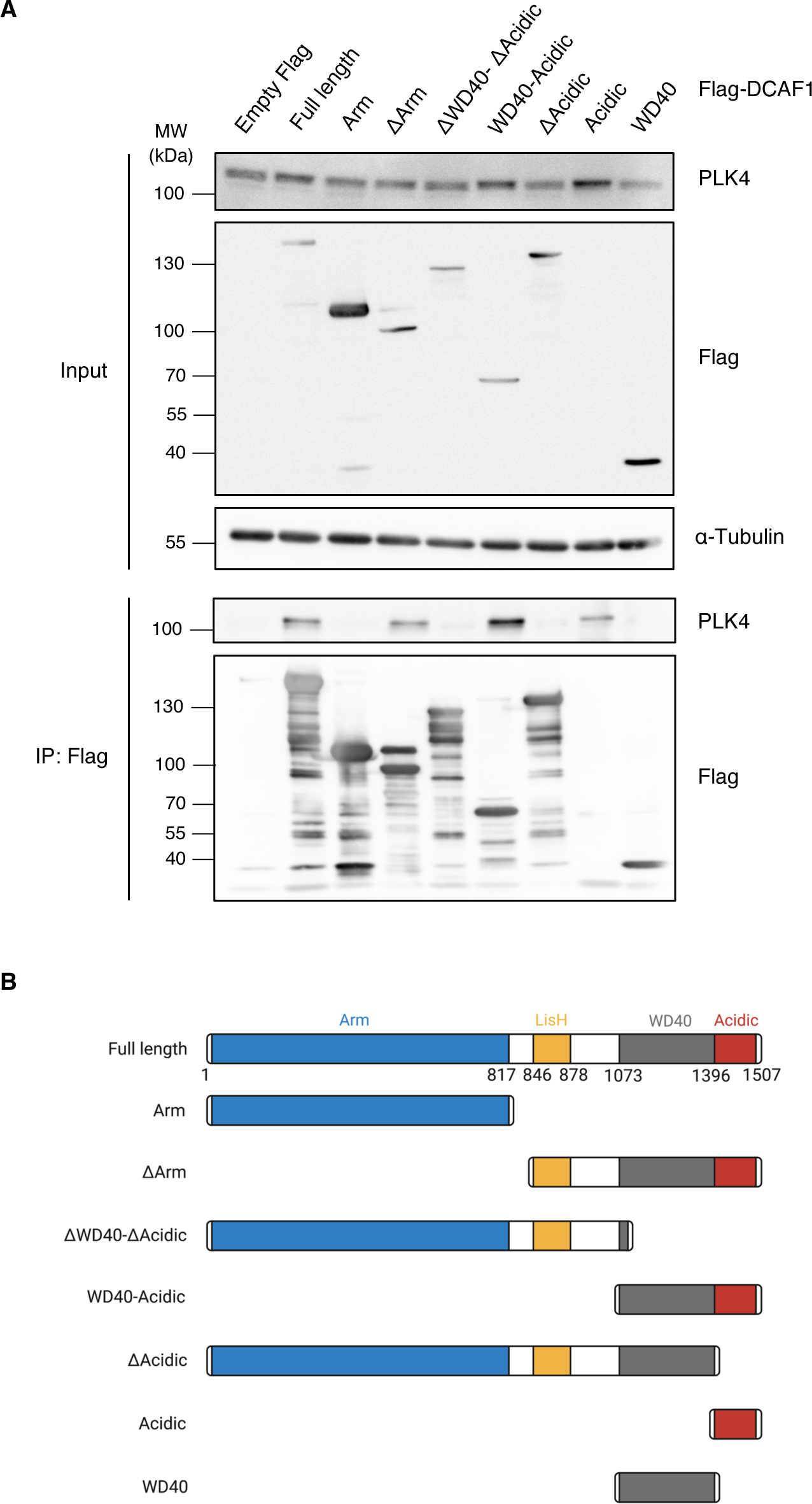
Flag-DCAF1 full length or different truncated fragments were overexpressed in HEK293T cells for 48 h. Co-precipitated PLK4 was detected by IP against the Flag tag and subsequent Western blot analysis. **(B)** Overview of the different Flag-DCAF1 fragments used in (A).

**Supplemental Figure 4.**
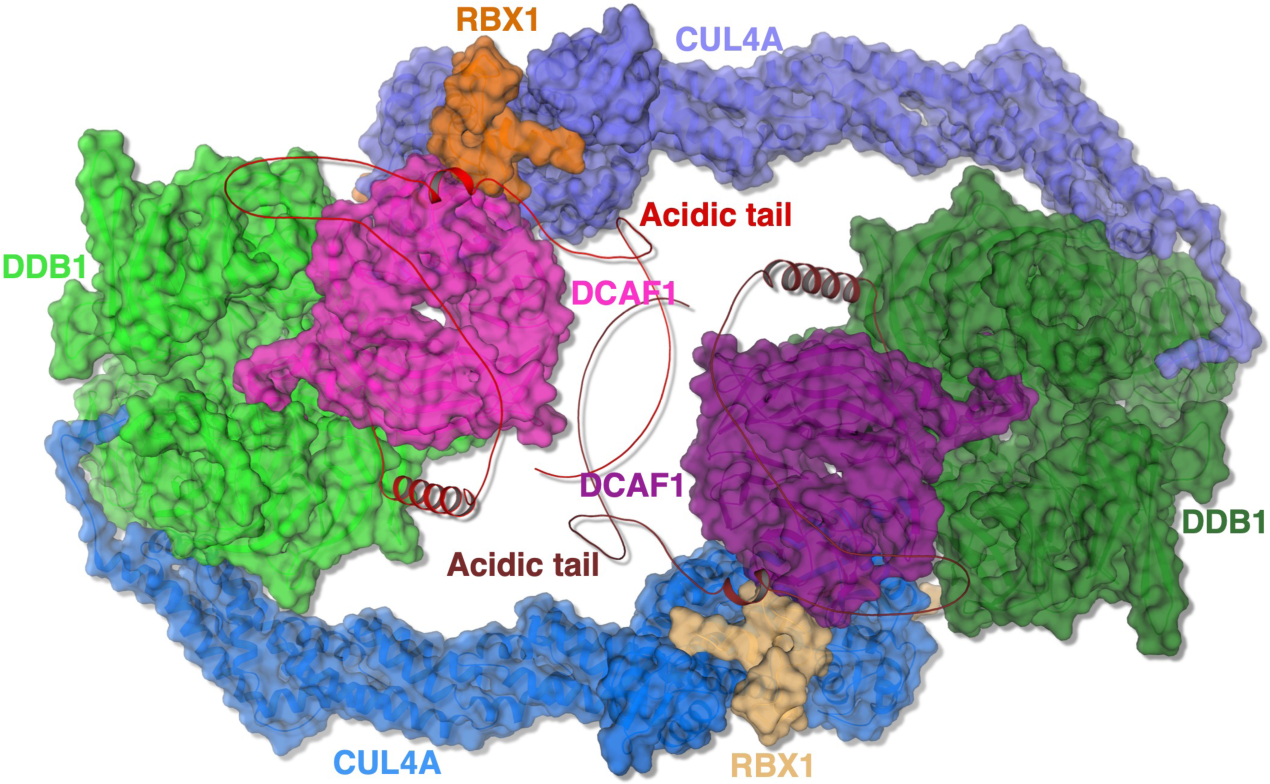
Cryo-EM structure of the CRL4^DCAF1^ complex and alphafold model of the DCAF1 acidic domain. The dimeric structure of DCAF1 (7OKQ) is shown and the names of protein components are indicated with the same color code. The unstructured acidic C-terminal tails of DCAF1 are shown in red and ruby.

**Supplemental Figure 5.**
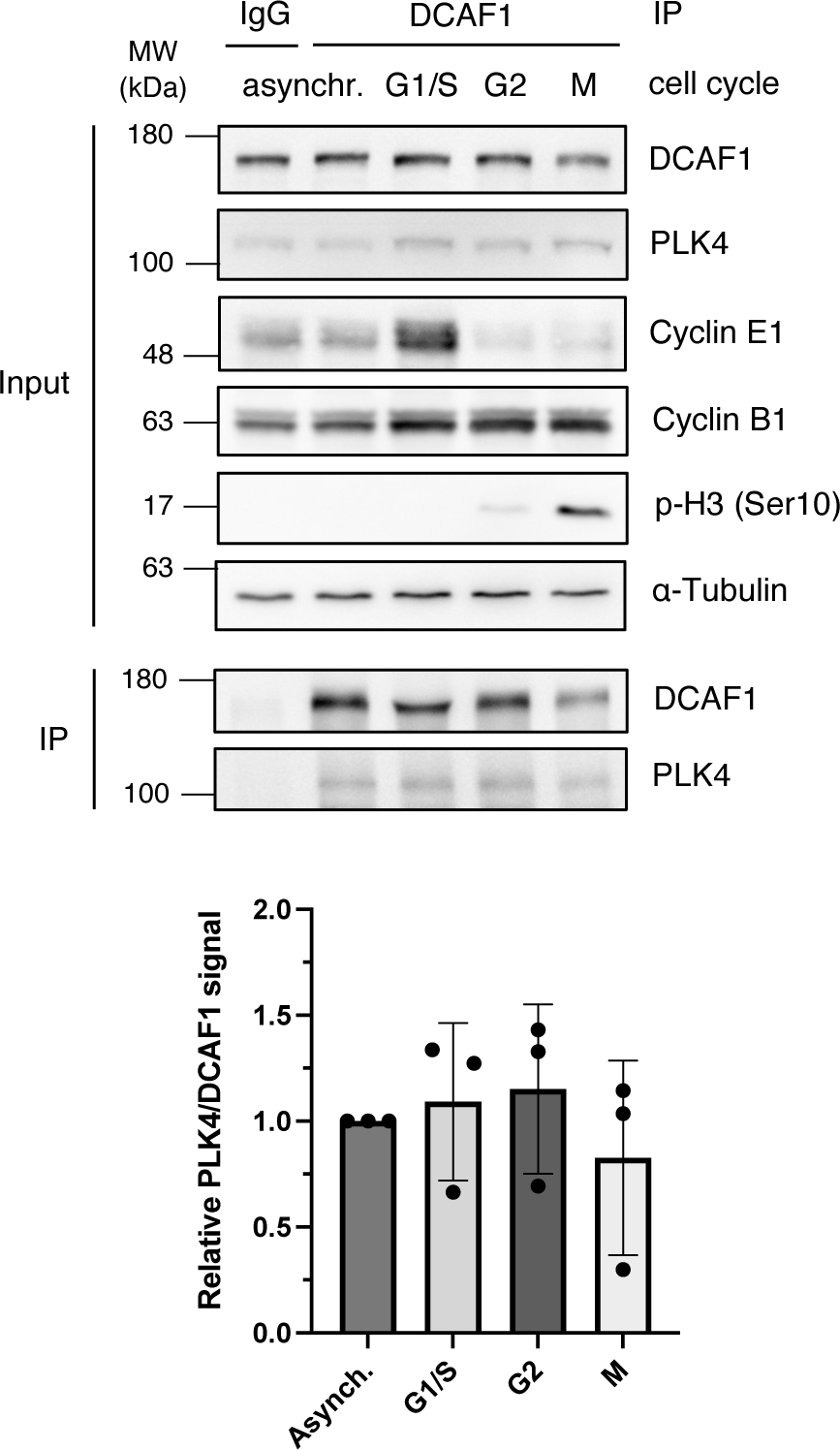
Cells were synchronized in G1/S phase by double thymidine arrest, in G2 phase by CDK1 inhibition with RO-3306 or in M phase by single thymidine and nocodazole arrest, as indicated. Endogenous DCAF1 was immunoprecipitated from cell lysates using unspecific IgG control or specific DCAF1 antibodies and protein G sepharose. Quantification of relative PLK4/ DCAF1 signal normalized to asynchronous cells. N = 3 independent experiments.

**Supplemental Figure 6.**
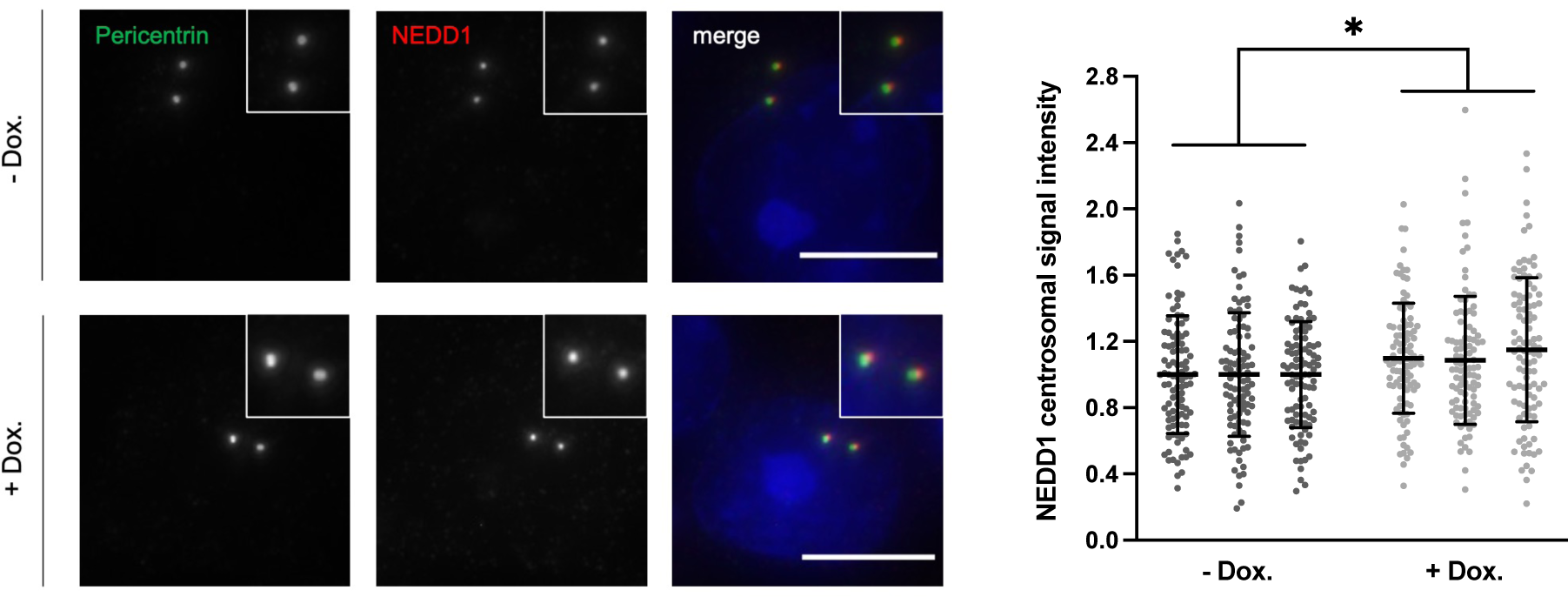
For knockdown of DCAF1, HeLa tet-on shDCAF1 cells were treated with 2 μg/ml doxycycline for 72 h prior to fixation. For immunofluorescence analysis, cells were stained with antibodies against pericentrin and NEDD1. Scale bar: 10 μm. Centrosomal signal intensities were quantified and normalized to the untreated control. * p < 0.05, unpaired Student’s t test with Welch’s correction, n = 3 independent experiments, 100 centrosomes analyzed per condition.

